# Distinct platelet interactions with soluble and immobilized von Willebrand factor modulate platelet adhesion and aggregation with differential impact on hemostasis and thrombosis

**DOI:** 10.1101/2024.01.31.578078

**Authors:** Yunfeng Chen, Zaverio M. Ruggeri

## Abstract

Arterial thrombosis is a prevailing and lethal pathological condition that remains difficult to treat or prevent without potentially serious side effects, mostly hemorrhagic in nature. Platelets and von Willebrand factor (VWF) have a recognized major role in the pathogenesis of arterial thrombosis. Platelets bind to surface immobilized VWF for initial adhesion to injured vascular sites, but also interact with soluble VWF to aggregate into thrombi, particularly under flow conditions creating elevated shear stress. Whether the binding of immobilized and soluble VWF to platelets is regulated by separate mechanisms and how they respectively regulate hemostasis and thrombosis remains unclear. Using targeted mutagenesis we engineered VWF to achieve modified binding kinetics with the platelet receptor glycoprotein (GP) Ibα and discovered that the interactions of immobilized and soluble VWF with platelets can be differentially regulated with distinct consequences on platelet adhesion and aggregation. Based on these results, we studied a monoclonal antibody, NMC4, known to bind to an epitope in the VWFA1 domain and to inhibit preferentially platelet aggregation under elevated shear stress conditions. We found that NMC4 was less efficient in reducing platelet adhesion to immobilized VWF than platelet aggregation mediated by soluble VWF and, surprisingly, also inhibited arterial thrombosis in a mouse model of ferric chloride-induced carotid artery occlusion at a dose that failed to prolong post-injury bleeding. Thus, our current findings help delineate interrelated biochemical and biophysical mechanisms underlying VWF function in vascular health; and suggest selective inhibition of VWF-mediated platelet aggregation as opposed to adhesion as a strategy to prevent arterial thrombosis while minimizing bleeding complications.

## Introduction

The balance between hemostasis and thrombosis is critical to human health. Diseases like hypertension, diabetes and obesity often leads to intensified arterial thrombosis in the coronary and cerebral circulations, which remains the leading cause of mortality and morbidity worldwide^1,2^. Distinct from hemostasis, arterial thrombosis is associated with elevated shear stress generated by stenosis, which has a prominent prothrombotic effect on platelets towards large-scale aggregation and vascular occlusion^3–6^. This phenomenon, termed ‘biomechanical platelet aggregation’, largely relies on the interaction between von Willebrand Factor (VWF) and platelet glycoprotein (GP) Ibα receptor^3,7^. Particularly, in pathologically high shear flow, biomechanical platelet aggregation can be solely achieved by VWF–GPIbα interaction in an integrin-independent fashion^8–10^. This indicates great potential of targeting VWF–GPIbα interaction for preventing arterial thrombosis^11–15^.

Unlike most other platelet ligands, VWF mediates both platelet adhesion and aggregation: in the immobilized form, VWF recruits circulating platelets and enables their adhesion onto the vascular wall, whereas the soluble form of VWF in the plasma mediates platelet-platelet cross-linking and therefore platelet aggregation. However, whether the activity of immobilized and soluble forms of VWF in platelet adhesion and aggregation is regulated by identical or distinctive mechanisms and how they respectively contribute to thrombosis and hemostasis are not fully understood. The VWF–GPIbα interaction is between VWFA1 domain and the GPIbα leucine rich repeats headpiece^16^. Although the majority of VWFA1 α4 and α5 helices is not in direct contact with GPIbα, they form a positively charged patch on VWFA1 surface (Fig. 1A), which can establish a long-range electrostatic attraction with a negatively charged patch on GPIbα^16^. This electrostatic attraction allows the positive charges in α4/α5 helices to facilitate the binding of platelet GPIbα to soluble VWF by directing the VWF molecules in solution to approach the GPIbα molecules, effectively increasing the rate of association. This is supported by previous works showing that removing certain positive charges from α4/α5 helices effectively inhibited soluble VWF binding to platelets^17^. The above binding mode is, by definition, three-dimensional (3D) binding, because at least on molecule is in the soluble form and thus has 3D movement freedom. However, the electrostatic attraction may become trivial for immobilized VWF-mediated platelet adhesion (by definition, 2D binding), where the accessibility of VWF to GPIbα is governed mainly by the proximity between the platelet surface and the VWF-immobilized surface during platelet “bumping and sweeping” driven by blood flow, and much less by molecular transport.

**Figure 1.**
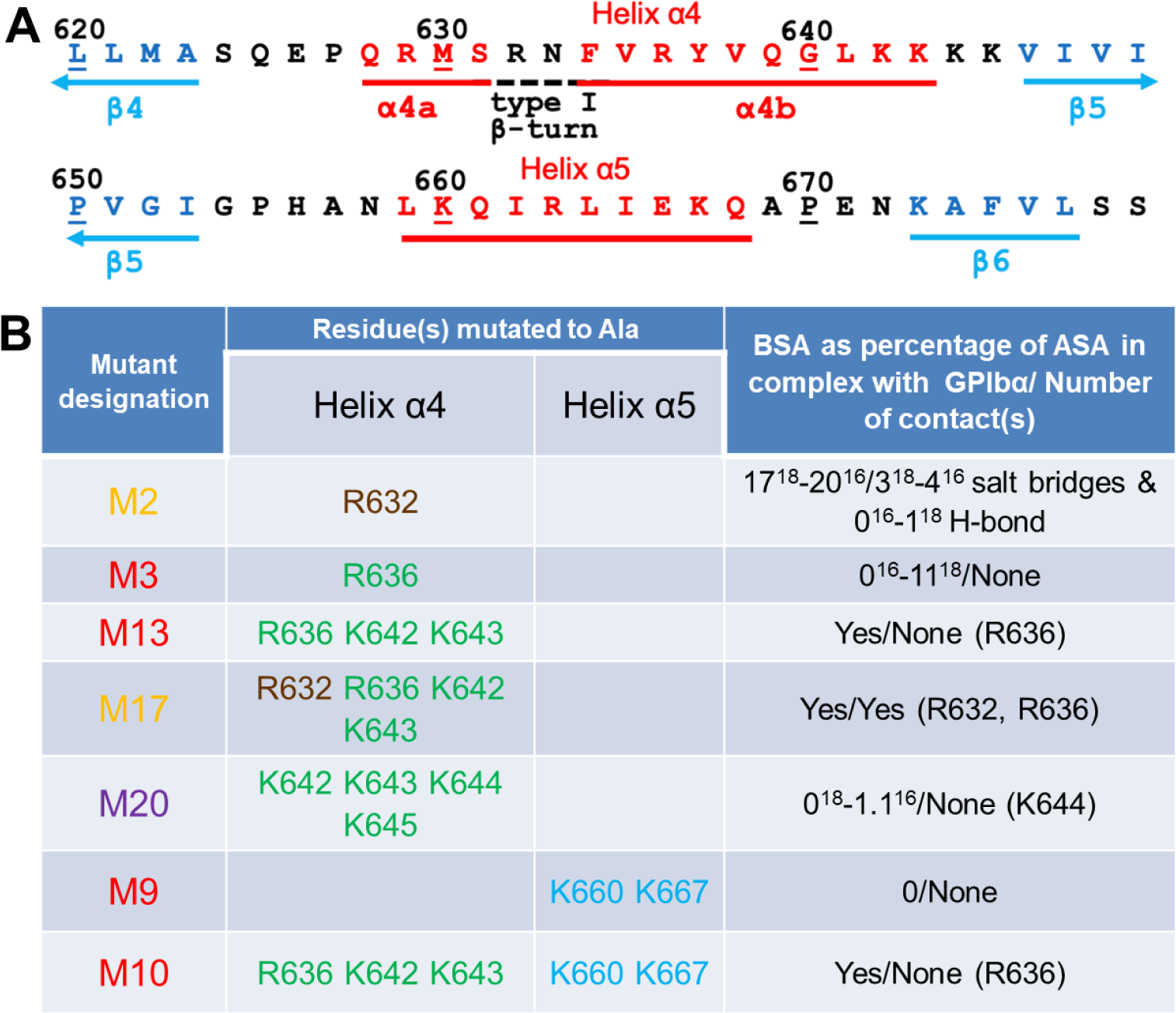
Sequence of VWFA1 α4/α5 helices and list of mutations investigated in this study. (**A**) Primary sequence and secondary structure of VWFA1 residues 620-679 highlighting α4/α5 helices (α-helices in red, β-sheets in blue, connecting loops and turns in black). (**B**) List of tested mutations that swap positive charged residues in VWFA1 α4/α5 helices with alanine. All amino acid position numbers are based on the sequence of mature VWF, which need to add 763 to derive the corresponding position numbers in pre-pro-VWF. R632 is in direct contact with GPIbα in the VWF–GPIbα binding complex, which is indicated in brown color. Two crystal structures respectively from the two indicated references are used to derive the BSA and contact information of the residues. BSA = Buried Surface Area; ASA = Accessible Surface Area; Contacts are bonds with interatomic distance <3.2 Å; data referred to the indicated amino acid residue.

Guided by the above rationale, we designed VWFA1 mutations in α4/α5 helices that inhibit VWF–GPIbα electrostatic interaction to differentially regulate the activity of VWF in its soluble and immobilized form. Agreeing with our hypothesis, removing GPIbα-non-contacting positive residues selectively inhibits soluble VWF in mediating biomechanical platelet aggregation, the extent of which is positively correlated with the number of positive charged removed. However, all these mutations maintain strong capacity of immobilized VWF in mediating platelet adhesion. Aligned with the effects of these mutations, a monoclonal VWF antibody, NMC4, was found to inhibit biomechanical platelet aggregation more efficiently than platelet adhesion, which also inhibited arterial thrombosis without compromising hemostasis under a certain dose range. Overall, our study identified a ‘hotspot’ within VWFA1 domain that can differentially mediate VWF activity in mediating platelet adhesion and aggregation, and suggested that inhibiting VWF in platelet aggregation but not platelet adhesion can potentially serve as a novel therapeutic strategy against arterial thrombosis that is devoid of the side effect of excessive bleeding.

## Results

### Design of VWFA1 mutations with selectively modified binding kinetics

A total of 7 VWFA1 mutations in α4/α5 helices were designed under different principles (Fig. 1B). Firstly, M3, M9, M13 and M10 swap 1, 2, 3 and 5 GPIbα-non-contacting positively charged amino acids with a neutrally charged amino acid, alanine, respectively. Among the amino acids subjected to mutation, R636 weakly contributes to VWF–GPIbα binding interface without having direct contact with GPIbα, as reflected by its low buried surface area and the absence of salt bridge revealed in the crystal structure of VWF–GPIbα complex^16,18^. Secondly, M20 mutates four consecutive positively charged amino acids in α4 helix, which was shown to drastically inhibit soluble VWF binding to platelets^17^, and is also suspected to causes local structural rearrangements besides inhibiting the electrostatic attraction. Lastly, in order to inspect the functional differences between GPIbα-contacting and -non-contacting residues, M2 and M17 (which is the add-up of M2 and M13) both mutate a positively charged amino acid that is in direct contact with GPIbα, R632^16,18^.

### Removing positive charges from VWFA1 α4/α5 helices impedes soluble VWF activity in platelet aggregation

We used aggregometry assay to test the activity of the VWF mutants (MTs) in platelet aggregation. A dimeric form of the VWFA1 construct (dA1) was produced instead of the full-length VWF to preclude platelet crosslinking by integrin–VWF C4 domain interaction. Platelet suspension added with ristocetin and varying concentrations of wild-type (WT) and MT dA1s were subjected to continuous stirring. A type 2B VWD gain-of-function (GOF) MT, I546V^19,20^, and a type 2M loss-of-function (LOF) MT, G561S^21,22^, were included as controls, which respectively mediated significantly stronger and weaker platelet aggregation than WT, confirming the reliability of the assay (Fig. 2A). These results were further confirmed by analyzing the maximal rate of aggregation (Fig. 2B,C). When mediated by M3, M9, M13 and M10 dA1, platelets showed progressively lesser degrees of aggregation (Fig. 2A, 2D-G). On the other hand, while M2 mutation showed negligible effect, M20 and M17 mutations also strongly inhibited platelet aggregation (Fig. 2H-J). The overall activity ranking of these mutants clearly shows a quantitative correlation between the number of positive charges removed and the residue VWF activity (Fig. 2K). Notably, changing the experimental mode to varying ristocetin concentration while maintaining the same dA1 concentration rendered the identical ranking, even at low ristocetin concentration. This rules out the alternative possibility that the weaker performance of the MT dA1s was due to defective association of ristocetin (Supp. Fig. 1).

**Figure 2.**
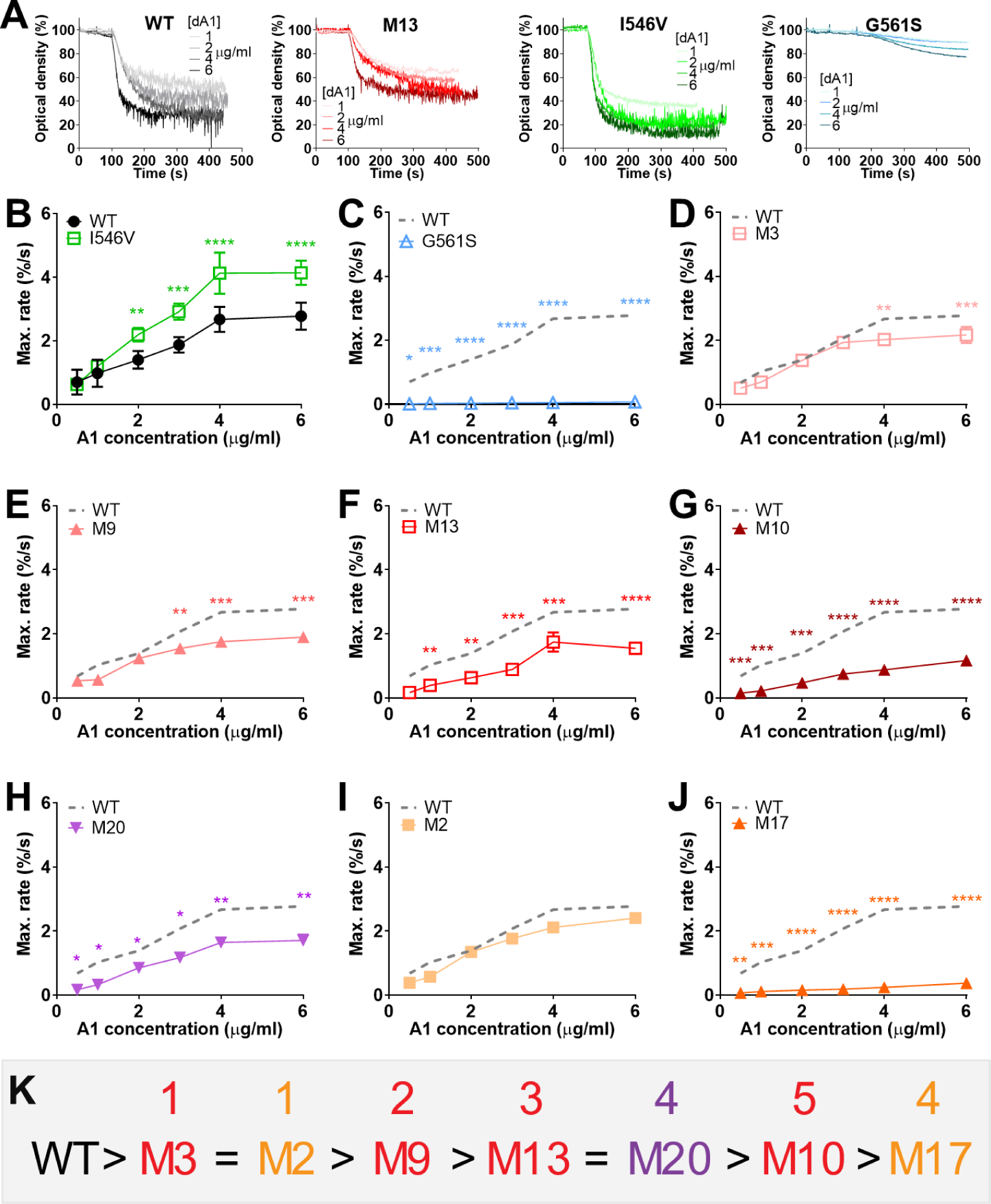
Removing positive charges from VWFA1 α4/α5 helices impedes VWF activity in platelet aggregation. (**A**) Representative optical density of platelet suspension vs. time curve in aggregometry assay with WT, M13, I546V and G561S dA1. The platelet suspension was adjusted to a platelet count of 200,000/μl, and added into a test tube with a stirring magnetic bar at t =0 s. dA1 was added to the suspension at t =40 s to reach the indicated concentrations, which was followed by the supplement of 1 mg/ml ristocetin after another 1 min. The optical density of the platelet suspension was monitored for a total of 450 s. (**B**-**J**) Mean±s.d. of maximal rate of platelet aggregation mediated by WT and I546V (B), G561S (C), M3 (D), M9 (E), M13 (F), M10 (G), M20 (H), M2 (I), M17 (J) dA1, respectively. In all the latter panels, the WT group was repeatedly shown as a gray dashed line for the convenience of comparison. *p < 0.05; **p < 0.01; ***p < 0.001; ****p < 0.0001 respectively, for comparison between WT and the mutants, assessed by two-way ANOVA, Sidak’s multiple comparisons test. (**K**) Rank of activity of WT and MT dA1s in mediating platelet aggregation. Numbers on top of the mutation names indicate the number of mutated residues.

### Removing positive charges from VWFA1 α4/α5 helices inhibits VWF–GPIbα 3D binding

Surface plasma resonance (SPR)^23^ is a technique that measures three-dimensional (3D) binding kinetics of molecular pairs, with one molecule immobilized and the other in the soluble form. Solutions of recombinant WT and MT monomeric VWFA1 (mA1) were perfused over an SPR chip coated with recombinant dimeric GPIbα headpiece (amino terminal). Agreeing with our aggregometry results, I546V and G561S mA1 respectively rendered significantly higher and lower affinities, higher and lower on-rates and lower and higher off-rates than WT (Fig. 3A-C). Along the removal of more positive charges, M3, M9, M13 and M10 mA1s showed a hierarchical decrease of affinity; on the other hand, while M2 mutation showed negligible effect, M20 and M17 mutations also strongly inhibited mA1–GPIbα 3D binding (Fig. 3A-C). Interestingly, the inhibitory effects of all these mutants are solely contributed by decreases in the on-rate, because their off-rates all remained comparable to WT. The overall affinity ranking of these mutants is highly consistent with the activity ranking derived from the aggregometry assay (Fig. 3D), altogether indicating that removing positive charges from VWFA1 α4/α5 helices inhibits VWF activity in the soluble form.

**Figure 3.**
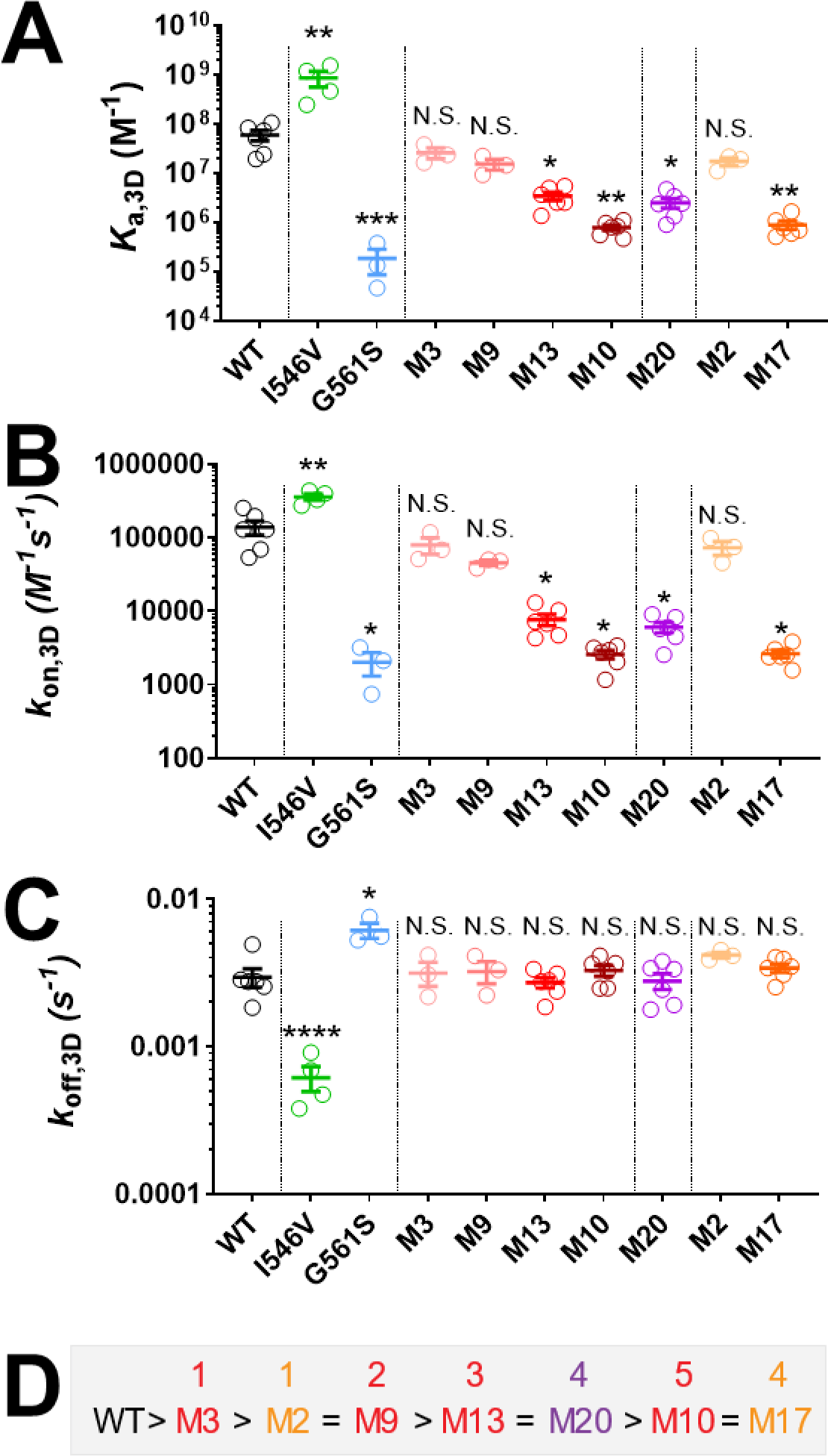
Removing positive charges from VWFA1 α4/α5 helices inhibits VWF–GPIbα 3D binding by reducing the one-rate. (**A**-**C**) Mean±s.d. (n ≥3) of 3D affinity (A), on-rate (B) and off-rate (C) of WT, I546V, G561S, M3, M9, M13, M10, M20, M2 and M17 mA1 binding to GPIbα headpiece measured by SPR. N.S. = not significant; *p < 0.05; **p < 0.01; ***p < 0.001; ****p < 0.0001 respectively, assessed by multiple t-tests. (**D**) Rank of activity of WT and MT mA1s in GPIbα headpiece binding in SPR. Numbers on top of the mutation names indicate the number of mutated residues.

### Removing GPIbα-non-contacting positive charges from VWFA1 α4/α5 helices reinforces immobilized VWF activity in platelet adhesion

We then used flow chamber assay to study how these mutations in VWFA1 α4/α5 helices affect platelet adhesion. WT and MT mA1s were respectively coated on coverslips with identical surface density, which was confirmed and calculated based on immunostaining (Supp. Fig. 2). Reconstituted human blood was then perfused over the surface. Under all shear rates tested (312-17,500 s^-1^) and with different mA1 coating densities, the surface coverage and rolling velocity of platelets on I546V mA1 were always respectively higher and lower than on WT, while no platelet adhesion was observed on G561S mA1 (Fig. 4A-C, 4K, Supp. Fig. 3A-C). These results confirm that the functional effects of I546V and G561S mutations in immobilized and soluble VWF are consistent (i.e., GOF and LOF, respectively). Interestingly, however, immobilized M3, M9, M13 and M10 VWFA1s all showed a GOF phenotype, manifested by high surface coverage (Fig. 4D-G, Supp. Fig. 3A,D,E) and slower rolling (Fig. 4L,M, Supp. Fig. 3B,H) of platelets, which is in sharp contrast to their LOF behaviors in the soluble form. These mutations are also in contrast to M20, which also only mutates GPIbα-non-contacting amino acids, but weakened platelet adhesion (Fig. 4H,N, Supp. Fig. 3F,I). The exceptional effect of M20 mutation is likely because the mutation of four consecutive lysines (K642-K645) results in local structural rearrangement and in turn disrupts VWF–GPIbα interaction via allosteric effects. On the other hand, although M2 mutation did not show substantial effects, its combination with M13 (i.e., M17) weakened instead of strengthened platelet adhesion, which is likely because mutating R632 directly disrupts VWF–GPIbα binding interface (Fig. 4I,J,O, Supp. Fig. 3F,I).

**Figure 4.**
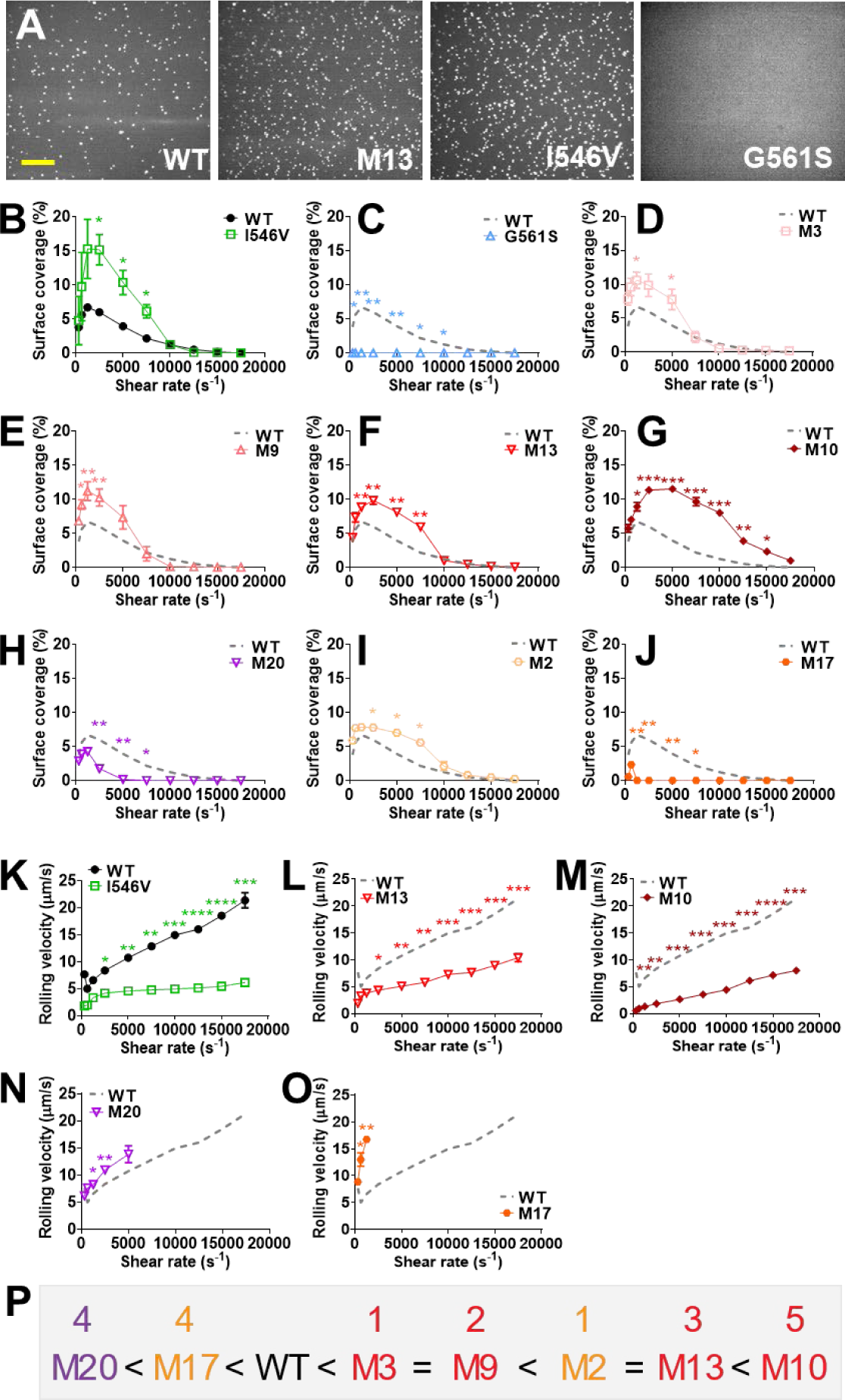
Removing positive charges from VWFA1 α4/α5 helices results in mixed effects in platelet adhesion. Reconstituted whole blood was perfused over a surface of 470/μm^2^ mA1 under 312-17500 s^-1^ shear rates. (**A**) Representative snapshots of platelets adhesion to surface coated with WT, M13, I546V or G561S mA1 under 1250 /s. Scale bar: 30 μm. (**B**-**J**) Mean±s.d. (n≥3) of platelet surface coverage on WT and I546V (B), G561S (C), M3 (D), M9 (E), M13 (F), M10 (G), M20 (H), M2 (I) and M17 (J) mA1, respectively. In all the latter panels, the WT group was repeatedly shown as a gray dashed line for the convenience of comparison. (**K**-**O**) Mean±s.d. (n≥3) of platelet rolling velocity on WT and I546V (K), M13 (L), M10 (M), M20 (N) and M17 (O) mA1, respectively. In all the latter panels, the WT group was repeatedly shown as a gray dashed line for the convenience of comparison. *p < 0.05; **p < 0.01; ***p < 0.001; ****p < 0.0001 respectively, for comparison between WT and the color-matched mutants, assessed by two-way ANOVA, Sidak’s multiple comparisons test. (**P**) Rank of activity of WT and MT mA1s in mediating platelet adhesion. Numbers on top of the mutation names indicate the number of mutated residues.

### Differential inhibitory effects of NMC4 in platelet adhesion vs. aggregation

NMC4 is a monoclonal antibody that inhibits VWFA1 binding to GPIbα^24,25^, and was shown to suppress thrombosis in transgenic mice expressing human GPIbα on platelets and human A1 in VWF (strain name: H1HA)^26^. Its docking interface with VWFA1 includes the respective association of three negatively charged Glu and Asp residues in NMC4 to three positively charged Arg residues in VWFA1 α4 helix, including R632 and R636^27^. Therefore, NMC4 likely acts as a mimetic of the mutations studied in this work by shielding part of VWFA1’s electrostatic interaction with GPIbα. To test this, aggregometry assay and flow chamber assay were respectively performed in the presence of varied concentrations of NMC4. Both assays showed a dose-dependent inhibitory effect of NMC4 (Fig. 5A,B). However, when the activities of dA1 (gauged by the maximal rate of platelet aggregation) and mA1 (gauged by platelet surface coverage) were normalized by their corresponding values under no NMC4 treatment, it was found that a much high concentration of NMC4 is required to reach the same level of inhibition in platelet adhesion as compared to platelet aggregation (Fig. 5C). These results indicate that NMC4 inhibits soluble VWF and its mediated platelet aggregation more efficiently than inhibiting immobilized VWF and its mediated platelet adhesion.

**Figure 5.**
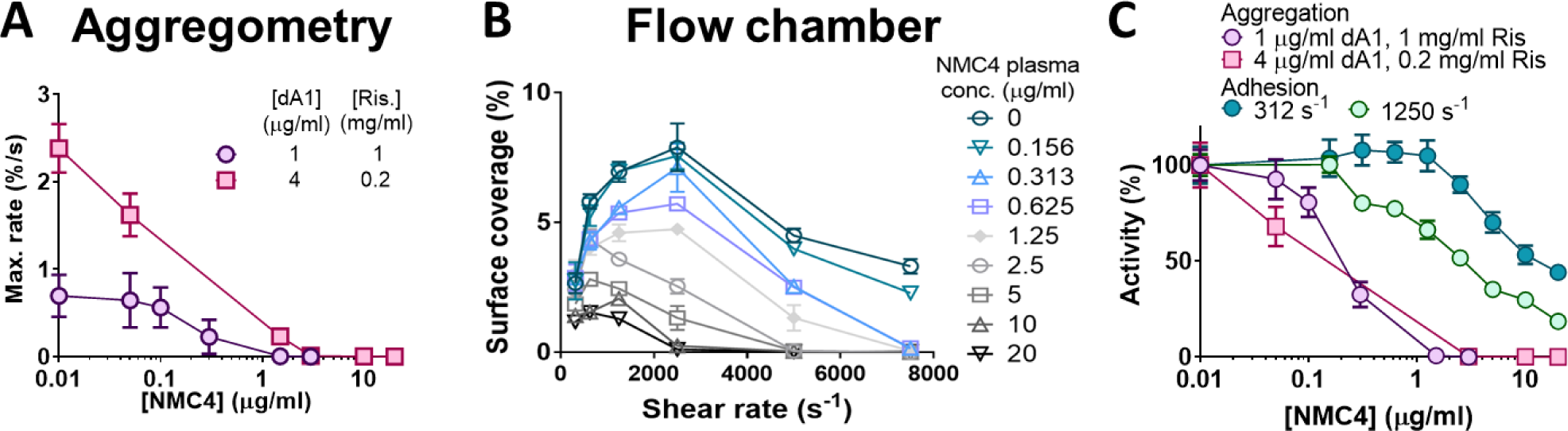
Differential inhibitory efficacy of NMC4 in platelet aggregation and adhesion. (**A**) Mean±s.d. of maximal rate of platelet aggregation mediated by dA1 in the presence of different concentrations of NMC4. (**B**) Mean±s.d. (n≥3), coverage of platelets on a surface coated with 470/μm^2^ monomeric VWFA1 in the presence of different concentrations of NMC4. (**C**) Dose-dependent activities of dA1 (A) and mA1 (B) (under 312 and 1250 s^-1^) were normalized by the values under no NMC4 treatment for direct comparison of inhibitory efficacy.

### NMC4 prevents arterial thrombosis without compromising hemostasis in a certain dose range

Finally, to understand the respective contributions of immobilized and soluble VWF to hemostasis and thrombosis, *in vivo* assays were performed using H1HA mice injected with different doses of NMC4. Injection of an equal volume of saline was used as negative control. In tail bleeding assay, at the dose of 2.5 μg (per 25 g mouse weight; applies to all *in vivo* doses) or higher, NMC4 significantly increased the bleeding time and blood loss, indicating compromised hemostasis; this inhibitory effect became statistically non-significant at the dose of 1.25 μg, and completely diminished at 0.625 μg (Fig. 6A). In contrast, in FeCl_3_-induced thrombosis assay, a NMC4 dose of 0.625 μg already showed considerable anti-thrombotic effects: the time to first and stable occlusion was substantially prolonged, and the flow index (volume flow rate of blood normalized by pre-surgery value) was also significantly reduced (Fig. 6B). Increasing the NMC4 dose to 1.25 μg further enhanced the anti-thrombotic effects, which completely prevented the formation of both unstable and stable thrombi and maintained flow index close to 1 in all tested mice (Fig. 6B). These results indicated that at the dose range of 0.625-1.25 μg per 25 g mouse weight, NMC4 effectively inhibits arterial thrombosis without significantly compromising hemostasis.

**Figure 6.**
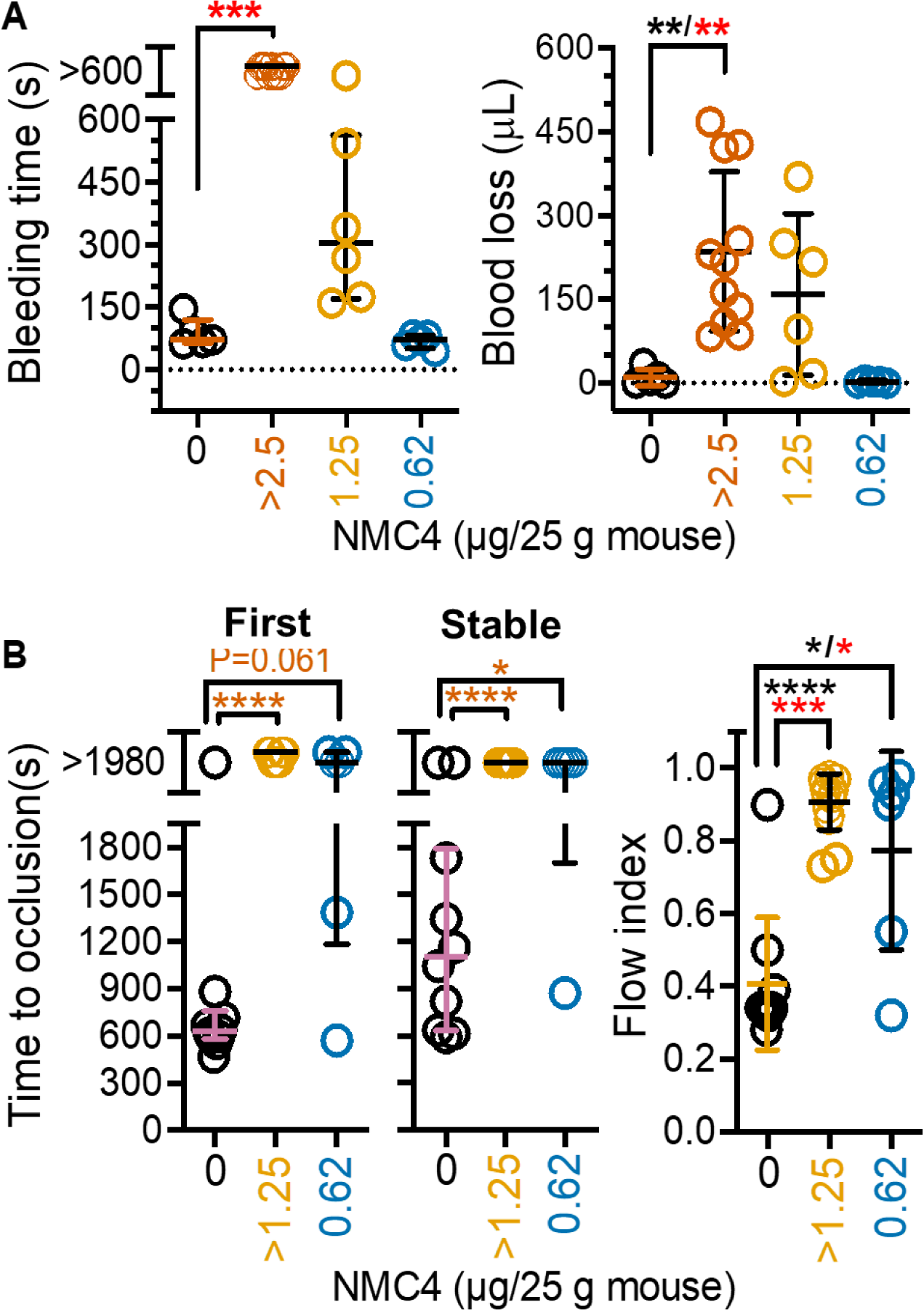
NMC4 dosed to have no effect on bleeding still reduces arterial occlusion in mice with humanized GPIbα and VWFA1. **(A)** Bleeding time and blood loss after tail clipping. The ≥2.5 μg group included 2 mice treated with 20 μg, 3 with 10, 1 with 5, and 5 with 2.5. **(B)** Carotid artery occlusion induced by 9% FeCl3·6H20. The ≥1.25 μg group included 9 mice treated with 20 and 6 with 1.25 μg. \**p* <0.05; \*\**p* <0.01; \*\*\**p* <0.001; \*\*\*\**p* <0.0001, assessed by nonparametric Kruskal-Wallis and Dunn’s post-test (red asterisks) or Brown-Forsythe-Welch ANOVA and Dunnett’s T3 post-test (black asterisks).

## Discussion

We discovered an atomic-level biophysical mechanism that differentially modulates the functions of immobilized and soluble VWF respectively in mediating platelet adhesion and platelet aggregation. Furthermore, we showed that preferentially inhibiting VWF-mediated platelet aggregation over platelet adhesion induces stronger anti-thrombotic effects than anti-hemostatic effects. Mechanistically, our findings can be summarized into the following model (Supp. Fig. 4): immobilized VWF mediates platelet adhesion via VWF–GPIbα 2D binding but not 3D binding, which plays a more important role in hemostasis than in thrombosis; in contrast, soluble VWF mediates platelet aggregation via VWF–GPIbα 3D binding, which contributes primarily to thrombosis but much less to hemostasis. Indeed, this theorem agrees with the fact that thrombosis heavily relies on platelet aggregation for endovascular blood clot growth, whereas platelet adhesion is required in the initial step of hemostasis to cover the vascular injury site. Another supporting evidence is an evolutionary insight from Schnaier *et al*, which pointed out that the avian thrombocytes have similar adhesion and signaling mechanisms as mammalian platelets, but are less efficient in aggregation; correspondingly, thrombocytes support hemostasis but cannot form occlusive thrombi^28^. The “biased” contributions of platelet aggregation and platelet adhesion to thrombosis and hemostasis is likely a general principle that also exist in other platelet receptor–ligand systems, the testing of which should inspire more safe anti-thrombotic strategies.

Despite previous evidences showing that VWF can support platelet aggregation independently under high shear laminar flow^8^ and that biomechanical platelet aggregation can be eliminated by an antibody inhibiting GPIbα^3^, the importance of VWF–GPIbα interaction-mediated platelet aggregation in the arterial system has always been underestimated. Instead, it is conventionally believed that VWF–GPIbα interaction primarily contributes to platelet adhesion, while platelet aggregation relies more on integrin α_IIb_β_3_. Here, the documented *in vitro* and *in vivo* effects of the anti-VWFA1 monoclonal antibody, NMC4, highlighted the importance of VWF–GPIbα interaction in platelet aggregation, which provides a paradigm shift to our understanding of VWF functions.

The binding of VWF to platelet GPIbα is regulated by multiple factors. Besides the binding interface itself, implicit regulating mechanisms also exist, such as the globular *vs*. elongated shape and the size of VWF multimer^29,30^ as well as VWFA1 auto-inhibition^31^. Here, our results indicate a previously underappreciated role of long-distance electrostatic attraction in VWF– GPIbα binding^32^ and underscored its physiological importance. They also provides an tentative explanation for certain von Willebrand diseases where mutating GPIbα-non-contacting amino acids in VWF causes profound impact to GPIbα binding^33^. Exactly why removing certain positively charged residues from the α4/α5 helices reinforces immobilized VWF function is still unclear, which could be possibly owing to an allosteric effect that makes the binding motif of VWF more accessible in the immobilized form^34–36^, or an orientational competition between VWF–GPIbα docking and electrostatic attraction. Nonetheless, our results on M3, M9, M13 and M10 provides the proof-of-concept for an unprecedented strategy that can achieve quantitatively controllable opposite effects on the affinities of immobilized and soluble forms of the same protein. The identification of VWFA1 α4/α5 helices as the regulatory ‘hotspot’ and the design strategy behind the mutations in this work should serve as a useful method in synthetic biology, which can be applied to protein design for reversely manipulating 3D and 2D binding kinetics.

Current antiplatelet agents (e.g., aspirin and clopidogrel) are inefficient in eliminating the prothrombotic effects of biomechanical platelet aggregation in high shear flow^3^ and severe stenosis^37–39^, due to the fact that they only inhibit platelet activation and aggregation mediated by soluble agonists (e.g., thromboxane A2 and ADP). Targeting VWF–GPIbα interaction and its mediated biomechanical platelet aggregation can potentially bridge this gap. However, due to the central role of VWF–GPIbα interaction in hemostasis, its blockage inevitably leads to excessive bleeding which has led to the failure of multiple clinical trials^40^. In this context, our finding that shielding the positively charged residues in VWFA1 α4/α5 helices preferentially inhibits arterial thrombosis over hemostasis is of clinical significance. The undesired effects of NMC4 in inhibiting platelet adhesion and prolonging bleeding under higher doses is likely because NMC4 targets certain VWFA1 residues that directly associate with GPIbα^27^, which might be alleviated by making NMC4 derivatives devoid of targeting GPIbα-contacting residues. Furthermore, our discoveries provide conceptual advances in the design of anti-thrombotics targeting VWF–GPIbα interaction, advocating for the development of ‘regulators’ instead of ‘blockers’ as a safer anti-thrombotic strategy.

## Materials and Methods

### Reagents and antibodies

Recombinant GPIbα headpiece (amino terminal, residues 2-290) and monomeric and dimeric VWFA1 (VWF subunit residues 445-733 fragments^8^ were expressed in *D. melanogaster* S2 cells and purified from culture supernatant^8,41^. The dimeric form of VWFA1 uses a thiol residue in the D3 domain to form a disulfide bond to link the two monomers^42^. NMC4 Fab fragment was produced as previously described^24^. Ristocetin was purchased from Chrono-log (Havertown, PA, USA). Borosilicate glass beads were from Distrilab Particle Technology (RC Leusden, The Netherlands).

### Transgenic mouse strain

All animal care and experimental procedures complied with the Guide for the Care and Use of Laboratory Animals, United States Department of Health and Human Services, and were approved by the Animal Care and Use Committee of The Scripps Research Institute. The generation of the H1HA mice was described in detail in a previous publication^26^.

### Surface Plasmon Resonance

To analyze the VWFA1-GPIbα interaction by surface plasmon resonance (SPR; Biacore 3000; GE Healthcare), human recombinant GPIbα headpiece followed by 133 residues of the SV40 large T antigen with Cys residues mediating dimerization was generated. A monoclonal antibody (MoAb) LJ-3A2 specific for the SV40 sequence was used for its capturing^41^. After linked to an SPR chip (HC200M; Xantec Bioanalytics), LJ-3A2 immobilized GPIb fragments in the appropriate orientation to measure kinetics parameters of VWFA1 binding. Varying concentrations (0.78-100 nM) of recombinant dimeric VWFA1 fragment with human or mouse sequence - VWF subunit residues 445-733 or 445-716, respectively (add 763 for the number in pre-pro-VWF) - were prepared in 135 mM NaCl, 20 mM Hepes (HBS, pH 7.4) containing 0.005% Tween 20. VWFA1 solutions were injected over the chip (association phase) at 75 μL/min for 3 min, followed by the injection of solution buffer for 20 min (dissociation phase).

Considering that VWFA1 domain can undergo dimeric self-association^27^, a two-phase association biophysical model was established to describe the binding of VWFA1 on immobilized GPIb and the subsequent binding of a second VWFA1. Similarly, a two-phase dissociation model was used to describe the dissociation of VWFA1 from the surface. By fitting the SPR curves to the two models, *k*_off,3D_ and *k*_on,3D_ were derived, which were then used to calculate the affinity *K*_a,3D_ (= *k*_on,3D_/*k*_off,3D_).

### Aggregometry assay

All procedures involving the collection of blood from healthy human donors were approved by Institutional Review Board of The Scripps Research Institute (protocol number IRB-12-5997). All human donor blood samples were obtained with informed consent. Blood was drawn from healthy donors into a syringe pre-loaded with buffered trisodium citrate debydrate 3.2% (0.11 M trisodium citrate, 20 mM citric acid, pH 5.2) at a 9:1 ratio. The blood was then centrifuged at 400 g for 15 min to derive the platelet rich plasma (PRP). The PRP was added with 1.2 U/mL apyrase (Sigma-aldrich, St. Louis, MO) and 10 μM Prostaglandin E_1_ (Cayman chemical company, Ann Arbor, MI) to inhibit platelet activation, and then centrifuged at 600 g for 15 min. The pellet was resuspended in Modified Tyrode buffer (MTB) pH 6.5 (135 mM NaCl, 11.9 mM NaHCO_3_, 2.9 mM KCl, 0.42 mM NaH_2_PO_4_, 10 mM Hepes, 5.5 mM dextrose) added with 1.2 U/mL apyrase and 10 μM Prostaglandin E_1_. The platelet suspension was centrifuged again at 600 g for 15 min, and finally resuspended in MTB pH 7.4, ready for experiments.

In each aggregometry assay, at time t =0 s, the platelet suspension was injected into a test tube with a stirring magnetic bar mounted on the aggregometry machine (Chrono-log, Havertown, PA). dA1 (and in some experiments NMC4) was added to the suspension at t =40 s to reach a preset final concentration, which was followed by the supplement of ristocetin after another 1 min. The final volume and platelet count of the suspension were respectively maintained at 250 µL and 200,000 /μL. The optical density of the platelet suspension was monitored for a total of 450 s, which was then normalized by the initial value and plotted against time. To derive the maximal rate of platelet aggregation in each test, we used a window of 10-point width to scan the “optical density vs. time” curve to derive the slope of each fragment. The absolute value of the steepest slope equals the maximal rate.

### Determining glass surface coating density of VWFA1

The bottom of a 96-well plate was removed and replaced by glass coverslips identical to those used for flow chamber assays via gluing. The wells were respectively coated with a series of concentrations (0-20 μg/ml) of monomeric VWFA1, with 3 repeats for each concentration. After 1 h, the plate was washed with HBS, pH 7.4, blocked with 5% bovine serum albumin (BSA; Sigma-Aldrich, St. Louis, MO) solution for 1 h, washed again, stained by 8 μg/mL NMC4-FITC solution for 30 min, washed again, and added with 50 μL HBS, pH 7.4. The fluorescence signal of each well was read by a plate reader (ThermoMax, Molecular Devices, San Jose, CA), which was converted to coating density values (number of molecules /μm^2^) using a calibration function.

To derive the calibration function, the fluorescence signals of 50 μL NMC4-FITC solutions with concentrations ranging from 0 to 20 μg/mL was measured by the same plate reader and plotted against the number of NMC4-FITC molecules subject to fluorescent excitation (N = 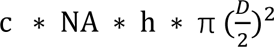, where *c* is the molar concentration of NMC4-FITC, *NA* is the Avogadro constant, and *h* and *D* respectively represent the height of the solution in the well and the laser beam diameter). The curve was then fitted with linear regression, and the fitting equation is the calibration function.

### Reconstituted whole blood flow chamber perfusion assay

Blood was drawn from healthy human donors into a syringe pre-loaded with ACD buffer (71 mM citric acid, 97 mM sodium citrate, 111 mM dextrose, pH 4.5) at a 5:1 ratio. The blood was then added with 10 μM Prostaglandin E_1_ and centrifuged at 400 g for 10 min. Platelet rich plasma was removed, and MTB pH 6.5 was added to maintain 40-45% hematocrit. An additional 10 μM of Prostaglandin E_1_ was added to prevent platelet activation. This procedure was repeated until the platelet count was lowered to ∼5,000 /μL. Afterwards, the blood was centrifuged at 1,500 g for 8 min, and the supernatant was removed and replaced with the same volume of MTB pH 7.4 added with 50 mg/mL BSA. Finally, the reconstituted blood was added with 10 μM Prostaglandin E_1_ and 10 μM DiOC_6_(3) (Life Technologies, Carlsbad, CA), and in some experiments NMC4 at different concentrations.

Perfusion experiments were conducted at 37°C by perfusing the reconstituted whole blood over a glass coverslip coated with mA1. The glass coverslips were assembled into a parallel flow chamber^43^. Driven by the aspiration of a syringe pump (Harvard Apparatus, Holliston, MA), blood was perfused through the chamber. The perfusion chamber was mounted on the stage of an inverted microscope (Axiovert 135M; Carl Zeiss, Thornwood, NY) for real-time visualization of platelet capturing on the immobilized substrate under different shear rates. Videos were analyzed by Fiji ImageJ 1.51u, and the platelet capturing was characterized by three parameters: surface coverage (percentage of the field of view covered by surface-captured platelets), number of new events (number of platelets newly captured to each unit area of the surface within each second) and rolling velocity.

### Tail bleeding assay

The tail bleeding assay was performed as described previously^26^. Mice were anesthetized using isoflurane in a precision vaporizer. The jugular vein was cannulated for injection of NMC4 10 minutes prior to injury. The tail was injured by clipping at 3 mm of the distal tip of the tail with a sterile scalpel blade and immersed in isotonic saline at 37°C. Blood collected from the tail was centrifuged and the cell pellet was resuspended in lysis buffer. The hemoglobin content was read with a plate reader at 450 nm, then converted to blood loss based on a calibration curve of known blood volumes. The time to complete cessation of blood flow was recorded as the bleeding time. After 600 seconds, persistent hemorrhage was stopped by cauterizing the tail wound.

### FeCl_3_-induced thrombosis assay

Mice were anesthetized using isoflurane in a precision vaporizer. The jugular vein was cannulated for injection of NMC4 10 minutes prior to injury. The common carotid artery of anesthetized mice was dissected, and an ultrasound flow probe was positioned around the vessel^26,44^. To facilitate the injury, a thin strip of silicone was placed beneath the artery between the probe and the chest. After measuring baseline flow, buffer was removed and the carotid artery was dried with filter paper. A 0.8-μL droplet of 9% (0.34 M) FeCl_3_ 6H_2_O was applied to the adventitia surface over the silicone strip for 3 minutes. Afterward, the surrounding area was washed with buffer resuming carotid blood flow monitoring which continued for 30 minutes (33 minutes total observation time). The artery was considered occluded when flow was <0.1 mL/min. Stable occlusion was defined as flow rate <0.1 mL/min for at least 10 minutes. A flow index was calculated as the ratio between the volume of blood that flowed through the carotid artery during the observation time and the volume calculated assuming constant baseline flow^8^.

### Statistical analysis

Statistical significance of the differences between two groups was determined by Student *t*-test or multiple t-test. For multi-group analysis, two-way ANOVA and Turkey test, nonparametric Kruskal-Wallis and Dunn’s post-test, or Brown-Forsythe-Welch ANOVA and Dunnett’s T3 post-test was used. *P* values <0.05 were considered significant.

## Acknowledgements

This work was supported by National Institutes of Health grants HL-117722 and HL-135290 (Z.M.R.) and HL-153678 (Y.C.), and AHA postdoctoral fellowship (Y.C.).

## Author contributions

Y.C. supervised the study, designed and performed experiments, analyzed data and wrote the paper; Z.M.R. supervised the study, designed experiments and wrote the paper. Research activities related to this work were complied with relevant ethical regulations.

