## Supplementary Figure 1-4 for "Distinct platelet interactions with soluble and immobilized von Willebrand factor modulate platelet adhesion and aggregation with differential impact on hemostasis and thrombosis"

**This file includes:**

Figs. S1 to S4

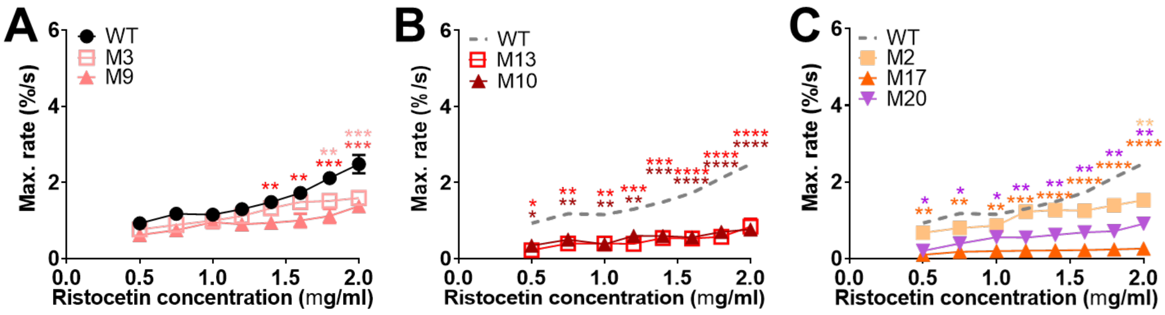


**Supplementary Figure 1. Testing mutant VWF activity in platelet aggregation using aggregometry assay**. The platelet suspension was adjusted to a platelet count of 200,000/µl, and added into a test tube with a stirring magnetic bar at t =0 s. 2 µg/ml dA1 was added to the suspension at t =40 s, which was followed by the supplement of ristocetin at the indicated concentrations after another 1 min. (**A-C**) Mean±s.d. of maximal rate of platelet aggregation mediated by WT, M3, M9 (A), M13, M10 (B), M2, M17 and M20 (C) dA1, respectively. In all the latter panels, the WT group was repeatedly shown as a gray dashed line for the convenience of comparison. *$p$ < 0.05; **$p$ < 0.01; ***$p$ < 0.001; ****$p$ < 0.0001 respectively, for comparison between WT and the mutants, assessed by two-way ANOVA, Sidak's multiple comparisons test.

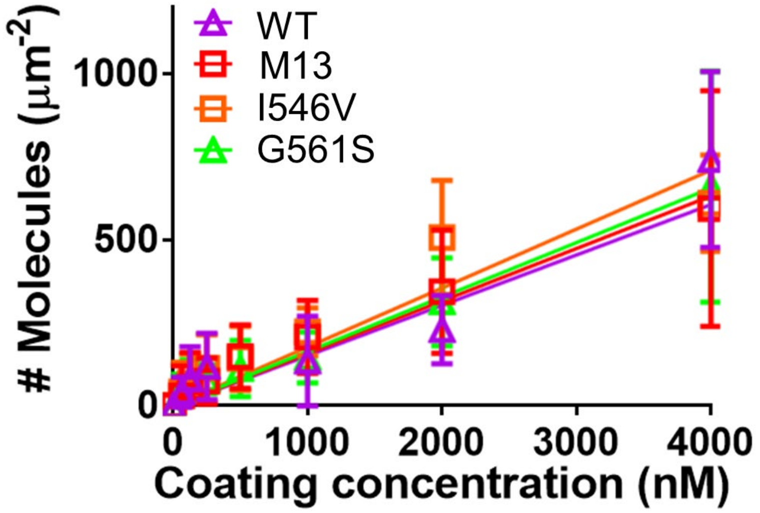


**Supplementary Figure 2. Mean±s.d. of surface coating density of WT, M13, I546V and G561S mA1 vs. coating solution concentration.** The data are fitted by linear regression passing through the original**.**

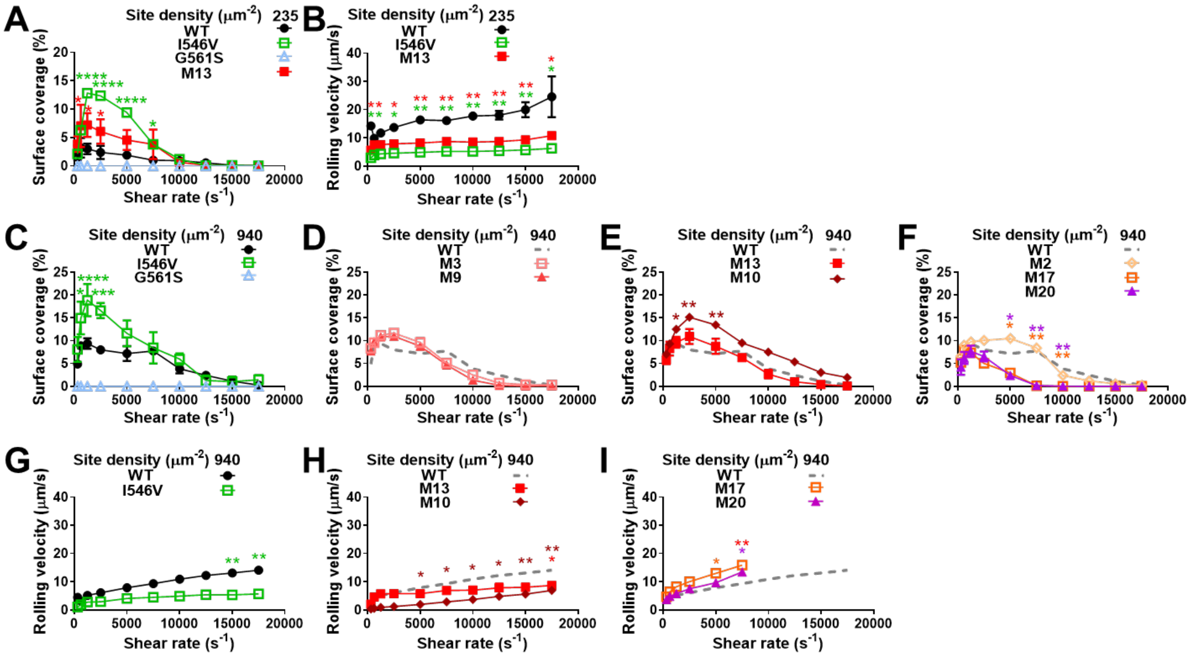

**Supplementary Figure 3. Assessing the activity of VWFA1 mutants in platelet adhesion under different coating conditions**. (**A**,**B**) Reconstituted whole blood was perfused over a surface of 235/μm$^2$ WT, I546V, G561S or M13 mA1 under 312-17500 s$^{-1}$ shear rates. Mean±s.d. (n≥3) of platelet surface coverage (A) and rolling velocity (B) were measured. (**C-I**) Reconstituted whole blood was perfused over a surface of 940/μm$^2$ mA1 under 312-17500 s$^{-1}$ shear rates. Mean±s.d. (n≥3) of platelet surface coverage on WT and I546V, G561S (C), M3, M9 (D), M13, M10 (E), M2, M17, M20 (F) mA1 was measured, respectively. In all the latter panels, the WT group was repeatedly shown as a gray dashed line for the convenience of comparison. Mean±s.d. (n≥3) of platelet rolling velocity on WT and I546V (G), M13, M10 (H), M17, M20 (I) mA1 was measured, respectively. In all the latter panels, the WT group was repeatedly shown as a gray dashed line for the convenience of comparison. *p < 0.05; **p < 0.01; ****p < 0.0001 respectively, for comparison between WT and the color-matched mutants, assessed by two-way ANOVA, Sidak's multiple comparisons test.

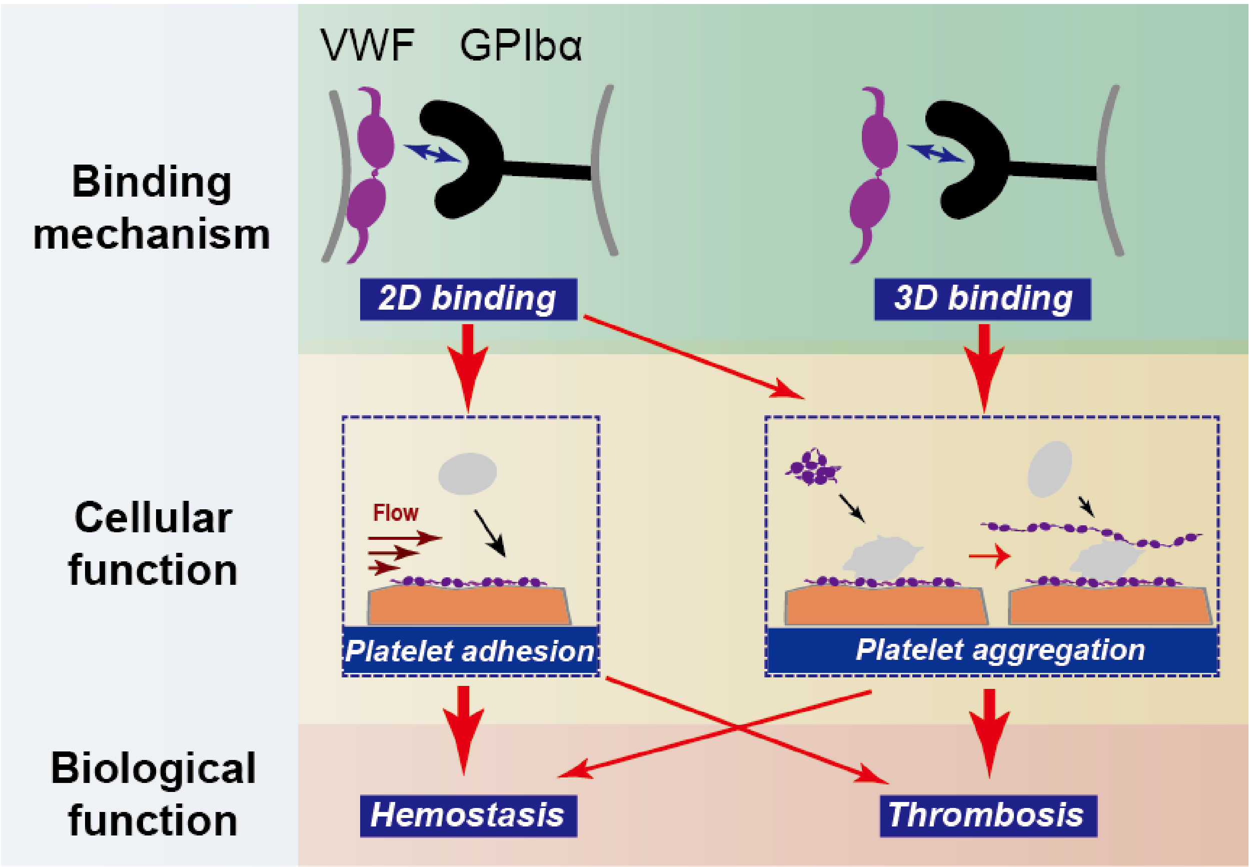


**Supplementary Figure 4. A model depicting the arterial functions of VWF regulated by different GPIbα-binding mechanisms.** Arrows indicate regulatory pathways of VWF–GPIbα binding mechanisms to the cellular functions of VWF, and of cellular functions to the biological functions. The thickness of the arrows reflects the relative importance of each pathway. VWF-mediated platelet adhesion is solely governed by 2D binding, which plays a primary role in hemostasis. In contrast, VWF-mediated platelet aggregation primarily contributes to thrombosis, which is mainly regulated by 3D binding.
